## Supplementary material for "A single inactivating amino acid change in the SARS-CoV-2 NSP3 Mac1 domain attenuates viral replication and pathogenesis *in vivo*"

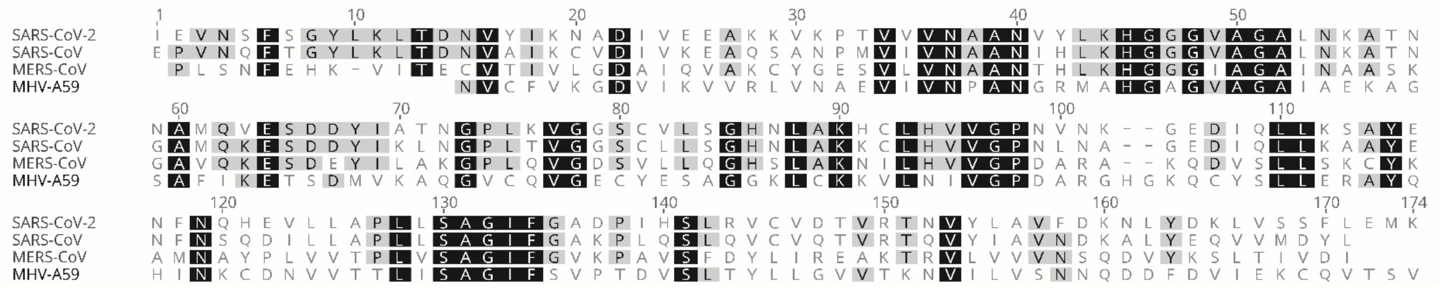

**Supplementary Figure 1**. Alignment of Mac1 sequences from several betacoronaviruses.
The Mac1 sequence of SARS-CoV-2, SARS-CoV, MERS-CoV, and MHV-A59 was obtained from uniport (P0DTC1, P0C6U8, K9N638, and P0C6V0, respectively) and aligned with Geneious Prime version 2022.2.1 using the Clustal Omega setting. Residues shaded in black are identical across all viruses, while residues shaded in grey are only partially conserved.

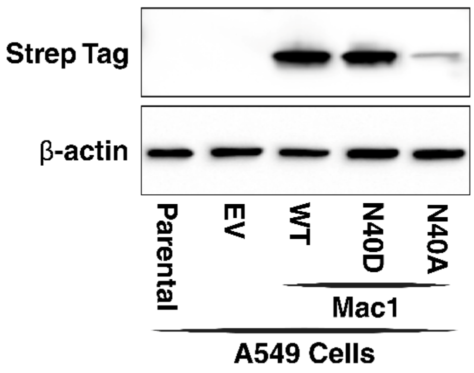


**Supplementary Figure 2.** Mac1$^{N40A}$ protein is less stable in cells than are Mac1$^{WT}$ or Mac1$^{N40D}$ proteins. Western blot analyses of cell lysates from A549 lung cancer cells lentivirally transduced with vectors expressing Strep-tagged Mac1$^{WT}$, Mac1$^{N40D}$ or Mac1$^{N40A}$ proteins. β-actin is shown as a loading control. The blot is representative of three independent replicates.

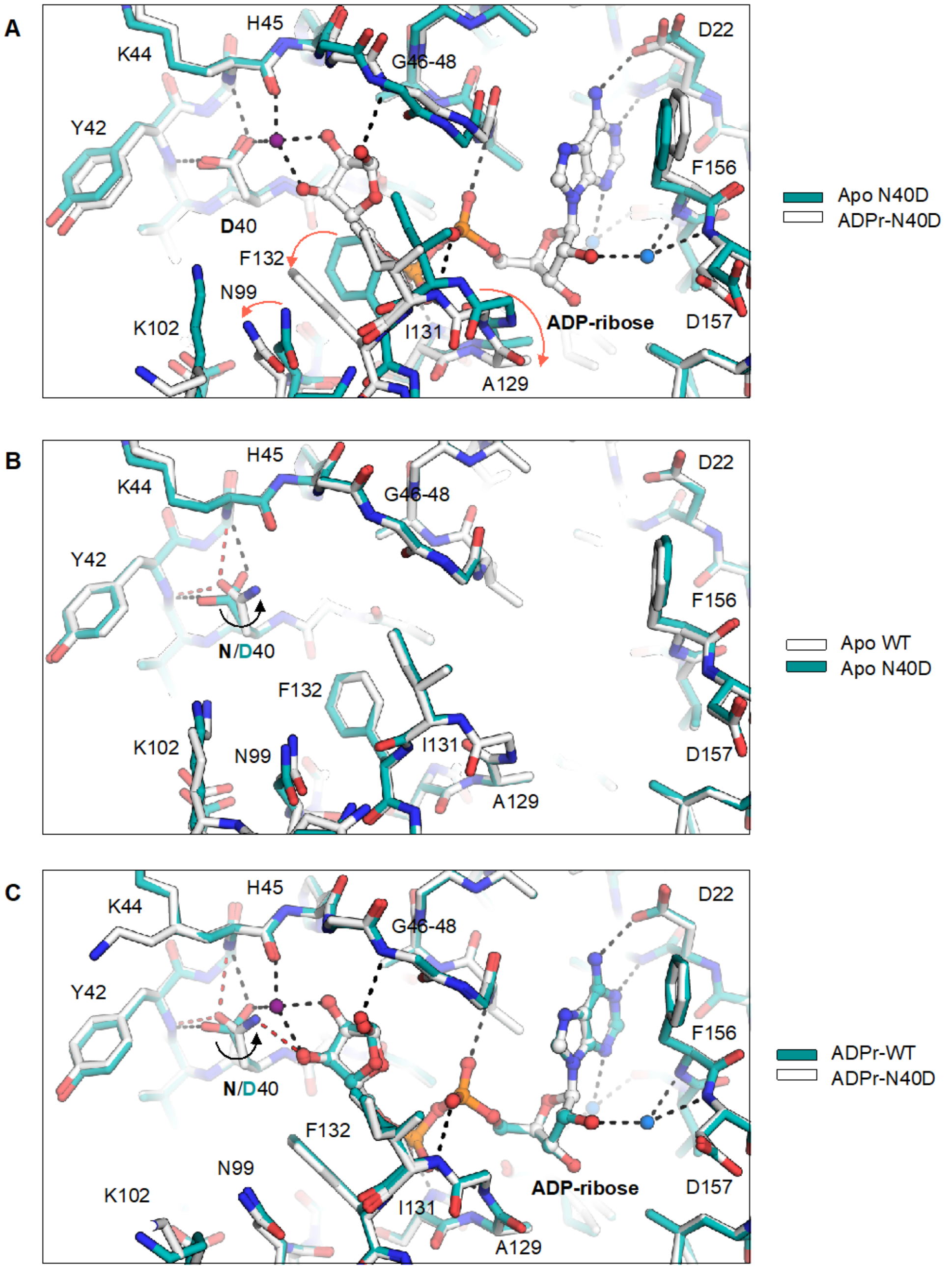


**Supplementary Figure 3.** Structure of Mac1 N40D determined by X-ray crystallography in apo- and ADP-ribose-bound state. (**A**) Alignment of apo- (PDB code 8SH6) and ADP-ribose-bound N40D (PDB code 8SH8) structures shows previously identified structural changes occurring upon ADP-ribose binding (e.g. Ala129 flip, Phe132 and Asn99 rotation, shown with orange arrows) (1). Selected hydrogen bonds between ADP-ribose and the active site are shown with dashed black lines. Three water molecules are shown that form bridging interactions between ADP-ribose and active site residues (purple/blue spheres). (**B**) Alignment of apo- WT (PDB code 7KQO) and N40D structures shows that there are only minor structural changes in the active site. The 80° rotation of the aspartic acid side chain relative to the asparagine side chain is indicated with a black arrow. (**C**) Alignment of ADP-ribose-bound WT (PDB code 7KQP) and N40D (PDB code 8SH8) shows that ADP-ribose binding is highly conserved despite the Asn40Asp mutation (RMSD across the 36 ADP-ribose atoms of 0.12 Å). The bridging water that is unique to N40D is shown with a purple sphere. Two conserved bridging waters are shown with blue spheres.

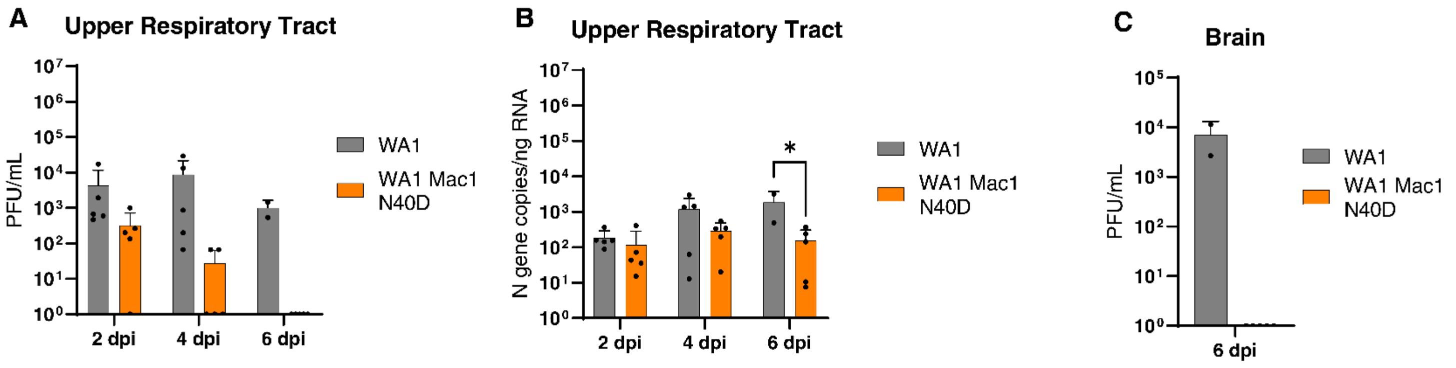


**Supplementary Figure 4**. SARS-CoV-2 Mac1 N40D mutant is attenuated in K18-hACE2 mice.

A) Viral particle abundance in the upper respiratory tract of mice infected with WT or Mac1 N40D mutant viruses was measured by plaque assay in VAT cells. Data is presented as mean +/- SD.

B) Viral RNA levels in upper respiratory tract were measured by RT-qPCR using a standard curve. Data is presented as mean +/- SD.

C) Viral particle abundance in the brains of mice infected with WT or Mac1 N40D mutant viruses at 6 days post infection was measured by plaque assay in VAT cells. Data is presented as mean +/- SD.

*, $p<0.05$ by two-sided Student's T-test.

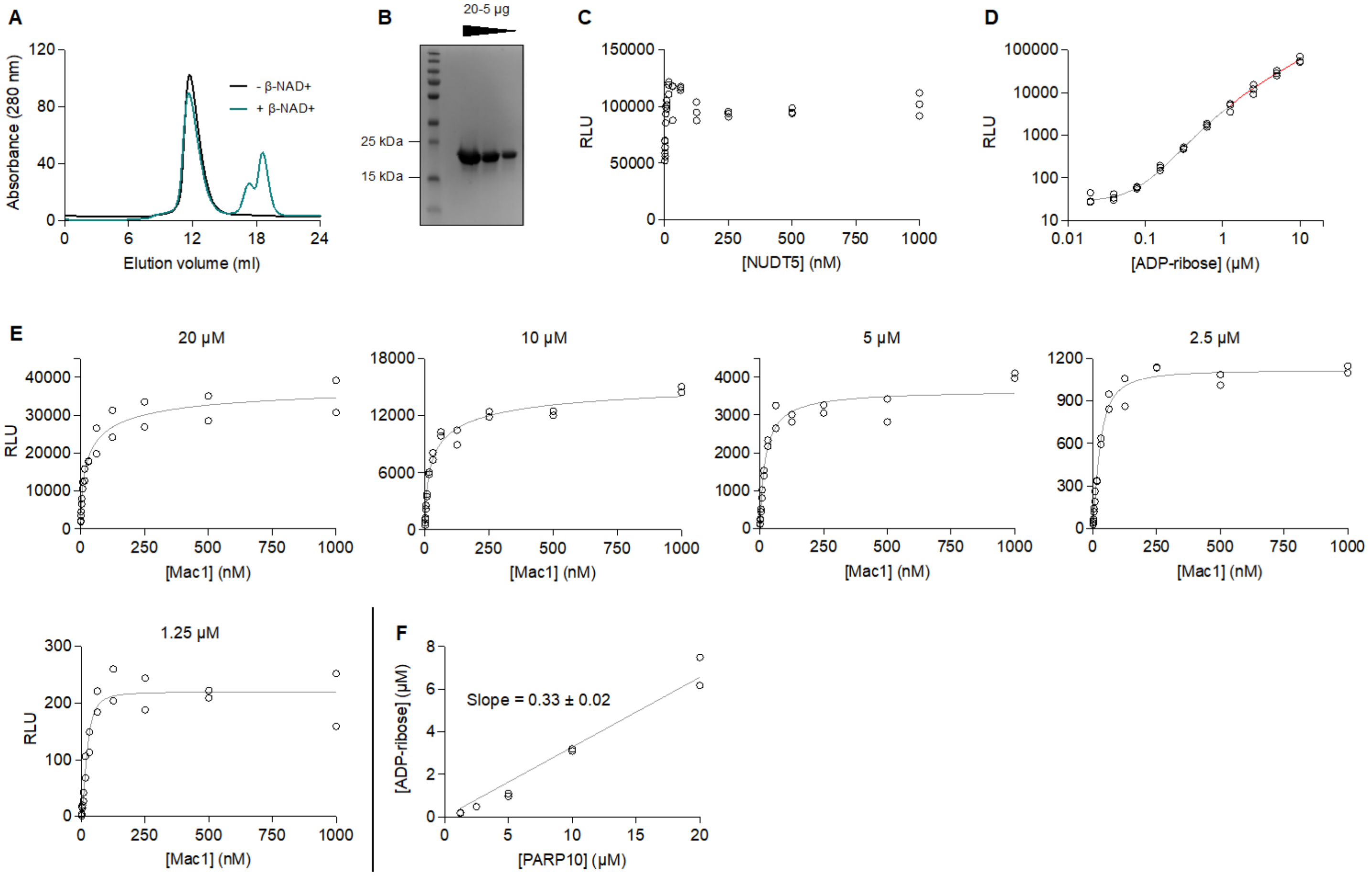


**Supplementary Figure 5**. Purification and quantification of MARylated PARP10. (**A**) Size exclusion chromatography (HiLoad 10/300 Superdex 75 pg column) of PARP10 with and without treatment with β-NAD+. The peaks eluting at ~18 mL are un-reacted β-NAD+ and/or hydrolyzed ADP-ribose. (**B**) SDS-PAGE (Biorad, 4569036) analysis of MARylated PARP10 with protein marker (ThermoFisher, 26616). Samples were boiled in SDS-PAGE loading dye for five minutes at 95°C and the gel was stained with Coomassie blue. (**C**) Titration of human NUDT5 phosphodiesterase with 20 µM ADP-ribose. AMP was detected using the AMP-Glo kit. Each condition was measured three times. (**D**) Log-log plot of the titration of ADP-ribose with 100 nM NUDT5. The highest three concentrations of ADP-ribose were fit using linear regression with GraphPad Prism (red line) and the remaining concentrations were fit to a four-parameter logistic equation using non-linear regression (gray line). Each condition was measured three times. (**E**) Titration of MARylated PARP10 and Mac1. Each plot shows the titration of Mac1 with a single concentration of MARylated PARP10. Each condition was measured twice, with the maximum luminescence determined by fitting a four-parameter logistic equation using non-linear regression (gray line). (**F**) ADP-ribose concentration generated by MARylated PARP10 hydrolysis plotted as a function of MARylated PARP10 concentration determined by absorbance at 280 nm. Data were fit by linear regression (gray line), with the line constrained to the X=0, Y=0 (the slope and standard error are indicated).

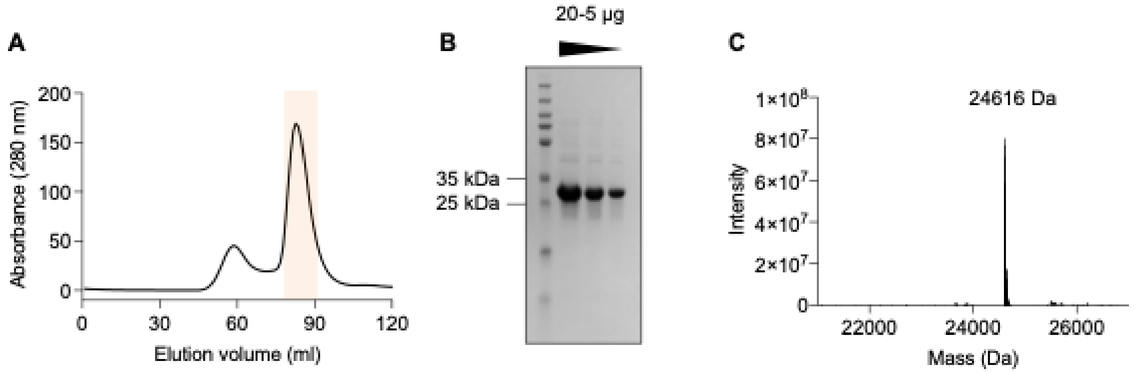


**Supplementary Figure 6**. Purification of human NUDT5. (**A**) Size exclusion chromatography (HiLoad 16/600 Superdex 200 pg column) of NUDT5 in the final purification step. The monomeric fractions pooled are shaded orange. (**B**) SDS-PAGE analysis of purified NUDT5 with protein marker (ThermoFisher, 26616). Samples were boiled in SDS-PAGE loading dye for five minutes at 95°C and the gel was stained with Coomassie blue. (**C**) Mass spectrum of purified NUDT5 deconvoluted with the maximum entropy method (MaxEnt1). The observed mass is within 3 Da of the mass calculated from the amino acid sequence (24619 Da).

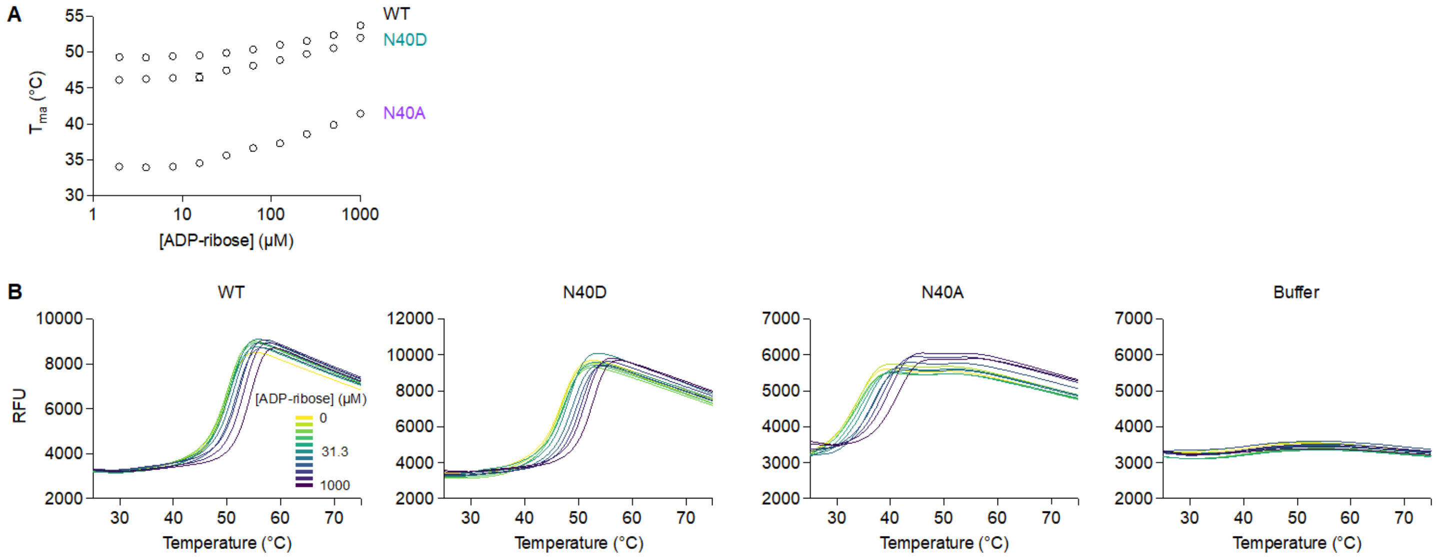


**Supplementary Figure 7**. Thermostability of WT, N40D and N40A mutants assessed by differential scanning fluorimetry with SYPRO orange. (**A**) Plot showing apparent melting temperatures ($T_{ma}$) for WT, N40D and N40A Mac1 as a function of ADP-ribose concentration. $T_{ma}$ values were calculated by DSFworld (2) using fitting model 1. Data is presented as the mean ± standard deviation for four technical replicates. (**B**) Raw fluorescence curves used to calculate $T_{ma}$ values. Data is presented as the mean for four technical replicates.

**Supplementary Table 1.** Data collection and refinement statistics for X-ray crystal structures reported in this work.

| | **Mac1 N40D** | **Mac1 N40D + ADP-ribose** |
|---|---|---|
| PDB code | 8SH6 | 8SH8 |
| **Experimental details** | | |
| Beamline | ALS 8.3.1 | ALS 8.3.1 |
| Temperature (K) | 100 | 100 |
| Transmission (%) | 100 | 100 |
| Wavelength (Å) | 0.7749 | 0.7749 |
| Energy (keV) | 16 | 16 |
| No. of images | 1800 | 1800 |
| Exposure time per image (s) | 0.1 | 0.2 |
| Total exposure time (s) | 180 | 360 |
| Beam size (μm) | 100x100 | 100x100 |
| Flux (photons/sec) | 8.60E+10 | 8.60E+10 |
| **Data reduction and refinement statistics** | | |
| Resolution range | 39.68-0.9 (0.91-0.9) | 39.39-0.9 (0.91-0.9) |
| Space group | P 43 | P 43 |
| Unit cell | 88.72 88.72 39.735 90 90 90 | 88.069 88.069 39.071 90 90 90 |
| Total reflections | 1496597 (41333) | 1454041 (40164) |
| Unique reflections | 227591 (7428) | 220438 (7138) |
| Multiplicity | 6.6 (5.6) | 6.6 (5.6) |
| Completeness (%) | 99.94 (98.72) | 99.88 (97.42) |
| Mean I/sigma(I) | 22.45 (1.91) | 19.26 (1.33) |
| Wilson B-factor | 7.76 | 7.83 |
| R-merge | 0.03619 (0.7356) | 0.04312 (1.084) |
| R-meas | 0.03931 (0.8125) | 0.0468 (1.195) |
| R-pim | 0.01522 (0.3385) | 0.01803 (0.4937) |
| CC1/2 | 1 (0.753) | 1 (0.602) |
| CC* | 1 (0.927) | 1 (0.867) |
| Reflections used in refinement | 227591 (7428) | 220438 (7138) |
| Reflections used for R-free | 11057 (372) | 10729 (346) |
| R-work | 0.1117 (0.2314) | 0.1143 (0.2625) |
| R-free | 0.1257 (0.2382) | 0.1233 (0.2976) |
| CC(work) | 0.976 (0.896) | 0.979 (0.844) |
| CC(free) | 0.973 (0.921) | 0.979 (0.834) |
| Number of non-hydrogen atoms | 3444 | 3322 |
| macromolecules | 2724 | 2772 |
| ligands | 0 | 36 |
| solvent | 720 | 514 |

| | | |
|---|---|---|
| Protein residues | 338 | 338 |
| RMS(bonds) | 0.004 | 0.019 |
| RMS(angles) | 0.82 | 0.85 |
| Ramachandran favored (%) | 100 | 99.1 |
| Ramachandran allowed (%) | 0 | 0.6 |
| Ramachandran outliers (%) | 0 | 0.3 |
| Rotamer outliers (%) | 0.99 | 0.32 |
| Clashscore | 3.1 | 3.71 |
| Average B-factor | 13.01 | 12.68 |
| macromolecules | 9.86 | 10.22 |
| ligands | | 7.05 |
| solvent | 24.93 | 26.35 |
